# CULTURE AND MAINTENANCE OF URINE-DERIVED, 3-DIMENSIONAL CANINE TRANSITIONAL CELL CARCINOMA ORGANOIDS

**DOI:** 10.1101/2021.08.24.457548

**Authors:** Savantha Thenuwara, Ben Schneider, Allison Mosichuk, Vojtech Gabriel, Christopher Zdyrski, Kimberly Dao, Chelsea Iennarella-Servantez, Madeline Colosimo, Dipak Sahoo, Agnes Bourgois-Mochel, Jean-Sebastien Palerme, Margaret Musser, Chad Johannes, Karin Allenspach, Jonathan P. Mochel

## Abstract

Bladder cancer is the ninth most common malignancy in the world. Transitional cell carcinoma (TCC), also referred to as urothelial carcinoma (UC) is the most common form of bladder cancer, occurring in 90% of cases. In this study, we explore urine-derived, 3-dimensional, canine TCC organoids as a possible model to study bladder TCC *ex vivo*. After establishing the cell lines, we subjected the 3D cells to RNA *in situ* hybridization (RNA-ISH) and cell viability assays. Overall, 3D cell culture from urine samples of TCC diagnosed canines expressed RNA biomarkers in a similar manner to parent tumors via RNA-ISH and showed more sensitivity to cisplatin treatment when compared to 2D human TCC cells. With further experimentation, canine TCC organoids could become an ideal model to study TCC *ex vivo*.

## INTRODUCTION

Bladder cancer is the ninth most common malignancy in the world (Ploeg et al., 2009). Over 81,000 new cases were diagnosed in the United States in 2020 alone, resulting in 17,980 deaths related to bladder cancer that year (Siegel et al., 2020). Bladder cancer is known to be a more common malignancy in men compared to women with 62,100 new cases in men and 19,300 new cases in women diagnosed in 2020 (Ploeg et al., 2009; Siegel et al., 2020). Urothelial carcinoma or transitional cell carcinoma (TCC) accounts for the majority (90%) of bladder cancer cases when categorized by histological type (Park et al., 2014). TCC can be further categorized into muscle-invasive (MIBC) and non-muscle invasive bladder cancer (NMIBC). NMIBC typically involves the mucosa, submucosa, or lamina propria of the bladder wall and accounts for 70% of newly diagnosed cases of bladder cancer (Abraham et al., 2012; Park et al., 2014). Treatment options for NMIBC involve intravesical chemotherapy or immunotherapy in conjunction with transurethral resection of bladder tumor (TURBT) to remove tumors or as diagnostic aids (Park et al., 2014; Kim and Patel, 2020). Approximately 25% of newly diagnosed cases of bladder cancer will be MIBC, which presents as a tumor that invades the muscular layer of the bladder wall (Abraham et al., 2012; Park et al., 2014). MIBC is associated with a higher rate of recurrence and poor overall prognosis compared to NMIBC (Abraham et al., 2012; Park et al., 2014). The standard course of treatment for MIBC is either medical management with cisplatin-based chemotherapy, or cisplatin as a neoadjuvant chemotherapy coupled with radical cystectomy (Aragon-Ching et al., 2018). However, even after radical cystectomy (removal of bladder), the 5-year survival rate for those with MIBC is about 40% (Stein et al., 2001). Additionally, developing targeted therapies for bladder cancer can be challenging due to the inherent heterogeneity seen in bladder cancer tumors (Thomsen et al., 2016, 2017).

Studying bladder cancer *ex vivo* presents its own challenges. While two-dimensional (2-D) cell culture organized in monolayers has been traditionally used in biomedical research, this system fails to capture the three dimensional (3-D) *in vivo* nature of tumor cells (Birgersdotter et al., 2005; Edmondson et al., 2014).

Attrition rates in oncology drug development are strikingly high, as only 5% of therapeutic leads that show anticancer activity in preclinical development are licensed after demonstrating sufficient efficacy in Phase III clinical testing (Hutchinson & Kirk, 2011). Although the underlying reasons for this high attrition are complex, they are to a large extent due to current suboptimal strategies for preclinical evaluation of therapeutic drug candidates (Wittenburg et al., 2011; Kumar et al., 2016; Nixon et al., 2017). In addition, a translational and predictive animal model of bladder cancer has been challenging to establish. Conventionally used mouse models require tumors to be experimentally induced as it is uncommon for rodents to develop tumors spontaneously (Arantes-Rodrigues et al., 2013). In essence, the phenotypic and molecular heterogeneity of MIBC tumors limits the translational value of conventional rodent models to evaluate therapeutic drug efficacy, since genetically modified mouse models do not effectively mimic the biological behavior of UC in humans (Ruan et al., 2019). For example, few genetically engineered mouse models display invasive phenotypes, which is an apparent limitation when modeling aggressive cancers like MIBC (Kobayashi et al., 2015). Also, the mouse urothelium is inherently refractory to developing cancer and mice do not develop metastases even when human tumors are implanted into immunodeficient animals (Kobayashi et al., 2015).

To address the issues presented above, 3D-cell culture has been presented as a viable model for studying bladder cancer, and particularly UC (Knapp et al., 2014; Vasyutin et al., 2019). 3D cell cultures of stem-cell-derived epithelial cells (termed organoids) are primary cells that grow into spheroids in a three-dimensional extracellular matrix formed by Matrigel^®^ or another similar matrix (Hughes et al., 2010; Fatehullah et al., 2016). Besides humans, similar descriptions of adult stem cells-derived 3D organoids have been presented in pigs, cats, and dogs (Mochel et al., 2017; Chandra et al., 2019; Ambrosini et al., 2020).

For bladder cancer organoids, the urothelium is the primary source of cells, as it is the predominant site of bladder cancer (Vasyutin et al., 2019). Studies have shown that these 3D cell cultures are better able to recreate the natural microenvironment of *in vivo* tumor cells in comparison to 2D cell cultures (Edmondson et al., 2014). Additionally, canines have been proposed as an attractive animal model for UC research (Knapp et al., 2014; Fulkerson et al., 2017; Minkler et al., 2021; Iennarella-Servantez et al., 2021; Gabriel et al., 2021). In fact, the vast majority (90%) of canine bladder cancer cases are muscle-invasive (Fulkerson et al., 2017). Also, there have been many noted similarities between canine and human UC cases. Cellular features, the heterogeneous nature of tumors, and patterns of metastasis are similar between species (Valli et al., 1995; Hughes et al., 2010; Knapp et al., 2015; Fulkerson et al., 2017). In addition, canine and human invasive bladder cancers respond similarly to chemotherapeutic treatment with cisplatin, carboplatin, and doxorubicin (Knapp et al., 2000; Boria et al., 2005; Robat et al., 2013; Fulkerson et al., 2017). Canine 3D organoids therefore have the potential to be a better model for studying UC *ex vivo*. Through this study, we aimed to explore culturing canine UC 3D-cells derived from urine samples as a non-invasive method of obtaining tumor cells for drug testing. In this report, we describe culturing canine UC organoids from samples obtained non-invasively through urine collection and performing RNA-ISH and Celltiter-Glo 3D Cell Viability Assay on organoids.

## METHODS

### Organoid and 2D cell culture maintenance

Urine samples were obtained from the Iowa State University Hixon-Lied Small Animal Hospital, and samples were kept on ice until cell isolation. Samples were processed no later than 4 hours after receiving them in the lab. Briefly, the urine sample was centrifuged at 700xg for 5 minutes, and the supernatant was removed. The remaining cell pellet was then incubated on ice for 20 minutes in Advanced DMEM/F12 (Gibco) + 1mg/mL Penstrep (PS). The DMEM/F12 + PS was spun down, removed, and the cell pellet was then washed with 1x complete chelating solution (CCS). The CCS was spun down and removed, and one final wash was done with DMEM/F12. Cells were then resuspended in Corning Matrigel (Corning, Cat no: 356231) and plated in 24 well plates with 30 μL of Matrigel per well. Organoids were then maintained as described in Ambrosini and colleagues (Ambrosini et al., 2020). With this procedure in place, a biobank of canine TCC organoids has been built, with organoids frozen in freezing media (50% Media, 40% FBS, 10% DMSO) and RNA*later™* (Invitrogen, Cat no: AM7021) preserved in liquid nitrogen, as well as fixed in formalin and embedded in paraffin.

2D cells were purchased from ATCC (ATCC, Cat no: CRL-2169). The cells, SW 780, were human TCC cells. Cells were plated in 75 cm^2^ tissue culture flasks using Leibovitz’s L-15 Medium (ATCC, Cat no: 30-2008) mixed with 10% FBS. These cells were passaged using TrypLE™ Express Enzyme (Thermofisher Scientific, Cat no: 12604013) once 80% confluency was reached. The cells used for the following experiments had only undergone one passage.

### RNAscope^®^

RNAscope^®^, an assay for tagging RNA in intact, fixed tissues via RNA *in situ* hybridization was performed on the organoids for phenotypic characterization (Wang et al., 2012). The protocol mentioned in Flanagan et al. (2012) was followed and four probes were used (Wang et al., 2012). FOXA1, CD44, KRT7 were experimental probes and Ubiquitin was used as a positive control. Probes were purchased from ACDBio, Inc. Newark, CA.

### Celltiter Glo 3D Cell Viability Assay

Celltiter-Glo 3D Cell Viability Assay (Promega, Cat no: G9681) was conducted on both 2D and 3D cell cultures to assess cytotoxicity in cell lines. Cells were passaged, counted, and split into opaque-walled 96-well plates, and 100μL of culture media was added to each well. Cells were then allowed to grow overnight. Cisplatin dissolved in 0.9% NaCl was added to cells (concentration intervals ranging from 0 – 3.5 μM) and the total volume in the well was brought up to 300μL. Cells were incubated with the drug for 24 or 48 hours. 200μL of media was removed from each well, and 100 μL of Celltiter-Glo reagent was added. Each well was pipetted 15 times in a manner that minimized bubbles and incubated for 25 minutes at room temperature. Luminescence was recorded. ATP standards (Promega, Cat no: P1132) were used in concentration intervals ranging from 25,000 nM to 10 nM and plated in the same plate as the cells. Results were analyzed using statistical software R (version 4.0.5).

The Celltiter-Glo 3D Cell Viability assay uses Luciferase to measure ATP. When the homogenous mix of Luciferase and Luciferin provided by the assay is added to the lysed cells, the Luciferin binds to ATP released from cells. This reaction, catalyzed by Luciferase, produces measurable luminescent signal that is proportional to the ATP present in the solution (Gerhardt et al., 1991).

## RESULTS

### 3D Organoid Cultures

Organoids were grown according to the method described above. **Figures 1.1A** and **1.2A** show light microscopy images taken soon after isolation, 3 days and 2 days respectively, and small spherical organoids along with cell debris and a brown color resulting from lysed red blood cells can be seen. Sample visibility increased 10 and 12 days after isolation showing larger, more robust organoids that are budding smaller, spherical organoids. At this point, TrypLE™ Express Enzyme was used to dissociate the organoids into smaller cell clusters during passaging. The before and after use of TrypLE™ Express can be seen in **Figures 1.3A** and **1.3B**. After multiple passages, organoids became almost “flower-like” in appearance, and even visible to the naked eye (**Figures 1.4A** and **1.4B**).

**Figure 1.**
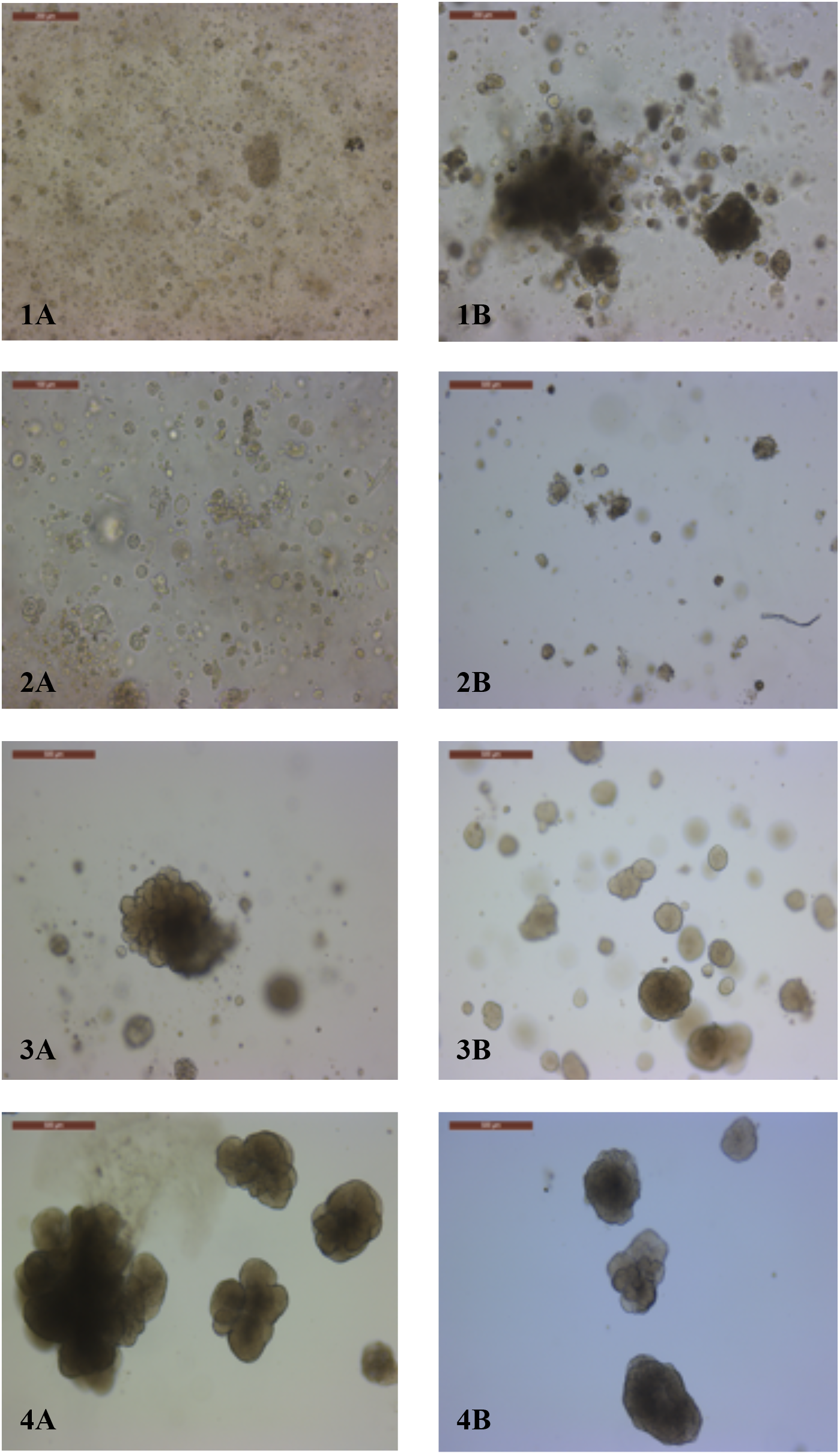
Light Microscopy images of urine-derived canine TCC organoids. Sample from canine “G” 3 days after isolation (1A) compared to 10 days after isolation (1B). Sample from canine “M” 2 days after isolation (2A) compared to 12 days after isolation (2B). Sample from canine “T” before 3(A) and after (3B) passaging with TrypLE™ Express. Sample from canine “T” (4A) 17 days post isolation, after 2 rounds of passaging and (4B) 15 days post isolation after 2 rounds of passaging.

### RNAscope^®^

Ubiquitin, used as positive control, showed staining for both organoid and tissue (not pictured herein). A transcriptional regulator, FOXA1 is expected to be downregulated in UC (DeGraff et al., 2012; Osei-Amponsa et al., 2020). Our RNAscope experiments show similar patterns of expression of CD44 between tumor tissue and tumor organoids acquired from the parent canine tumor (**Figures 2.1A** and **2.1B**). KRT7 is a member of the cytokeratin gene family and encodes type II cytokeratin, a relevant biomarker for detecting UC (Ichimi et al., 2009). As can be seen in **Figures 2.2A** and **2.2B**, in comparing tissue and organoid from the same parent tumor, we see similar levels of expression of KRT7. Knockout of FOXA1 in bladder cancer cells was associated with decreased E-cadherin expression and increased cell proliferation (DeGraff et al., 2012). As seen in **Figures 2.3A** and **2.3B**, FOXA1 is similarly poorly expressed in both tissue and organoid. CD44 is a cancer stem cell surface marker, and UC cells expressing CD44 have been shown to be more resistant to irradiation (Wu et al., 2017).

**Figure 2.**
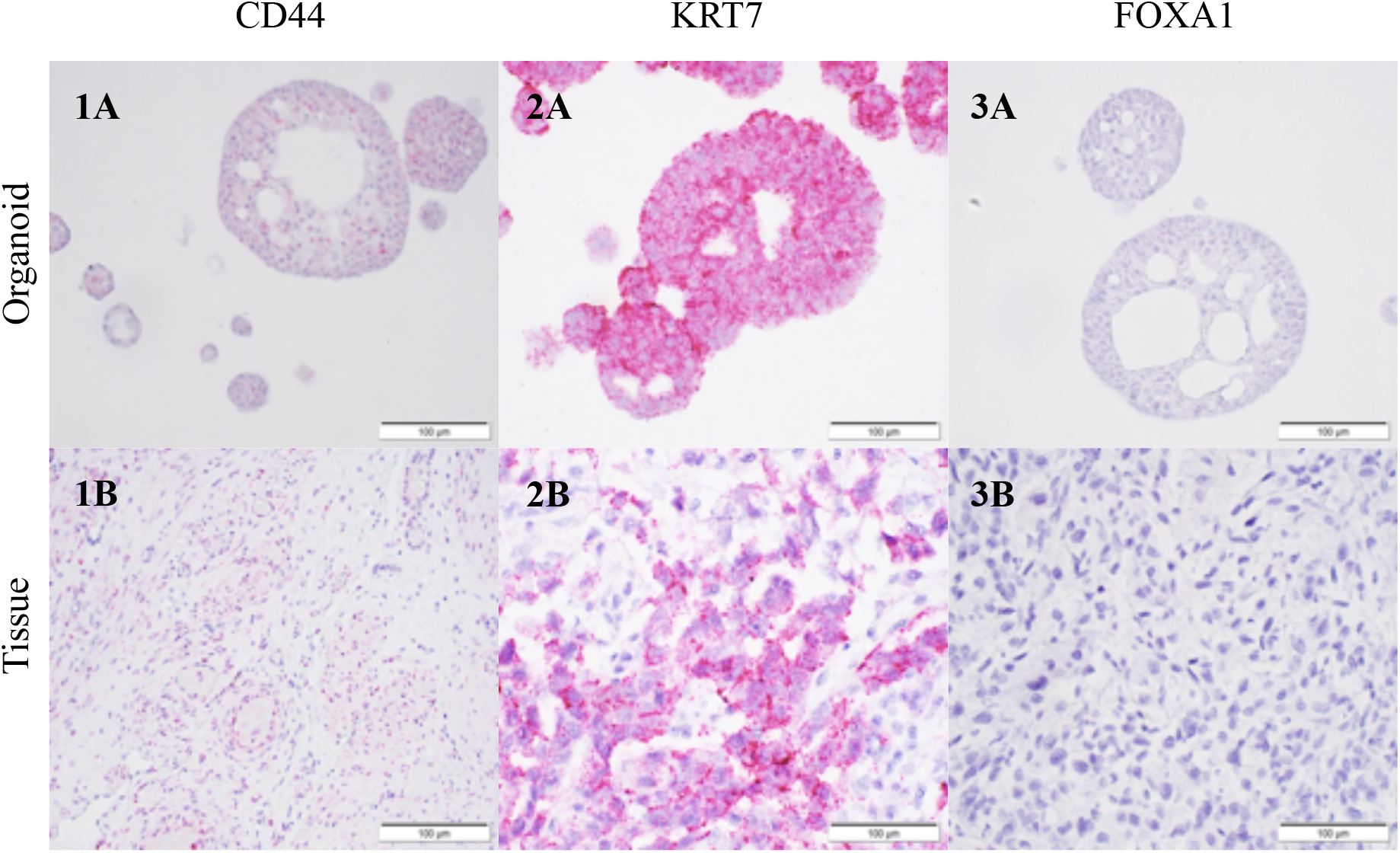
RNAscope^®^ staining on canine MIBC organoids and corresponding tumors. Organoid (1A) and tissue (1B) tagged with CD44 probes show similar patterns of CD44 expression. Organoid (2A) and tissue (2B) tagged with KRT7 show similar levels of expression of that RNA biomarker. Organoid (3A) and tissue (3B) tagged with FOXA1 show downregulation of FOXA1 in both tumor and organoid.

### Cell Viability

Using the packages “*Dose Finding*”, “*lattice*” and “*mvtnorm*” in statistical software R^1^, an exposure-response relationship was calculated between concentration of Cisplatin administered and concentration of ATP reported in nM, as seen in **Figure 3**. The X-axis of each graph shows the concentration intervals of Cisplatin (0μM, 0.25μM, 0.5μM, 1μM, 2μM, 3μM, 3.5μM) and Gemcitabine (10M, 1E-01M, 1E-03M, 1E-05M, 1E-07M, 1E-09M) Concentration ranges were chosen according to previously published work subjecting 2-dimensional urothelial carcinoma cells to the same drug. Depending on the shape of the exposure-response, an *Emax* or *Sigmoidal Emax* model was used to predict IC50 values and standard error (**Table 1**). The Y-axis displays measured concentrations of ATP in nM. Each point on the graph displays the mean ATP concentration of 8 replicates for the 2D data and of 3 replicates for the 3D data. Each graph also displays a 95% confidence interval.

**Figure 3.**
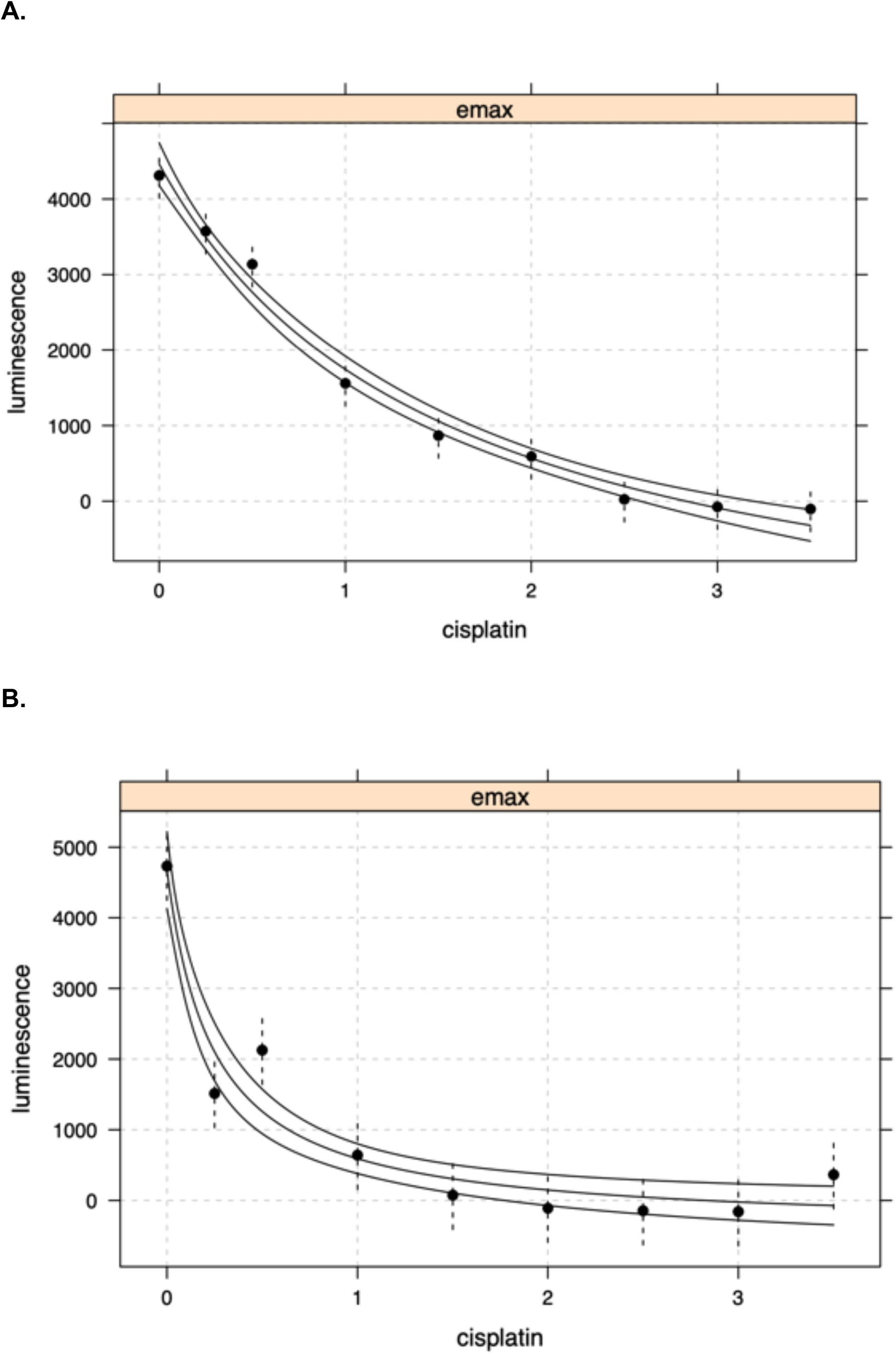
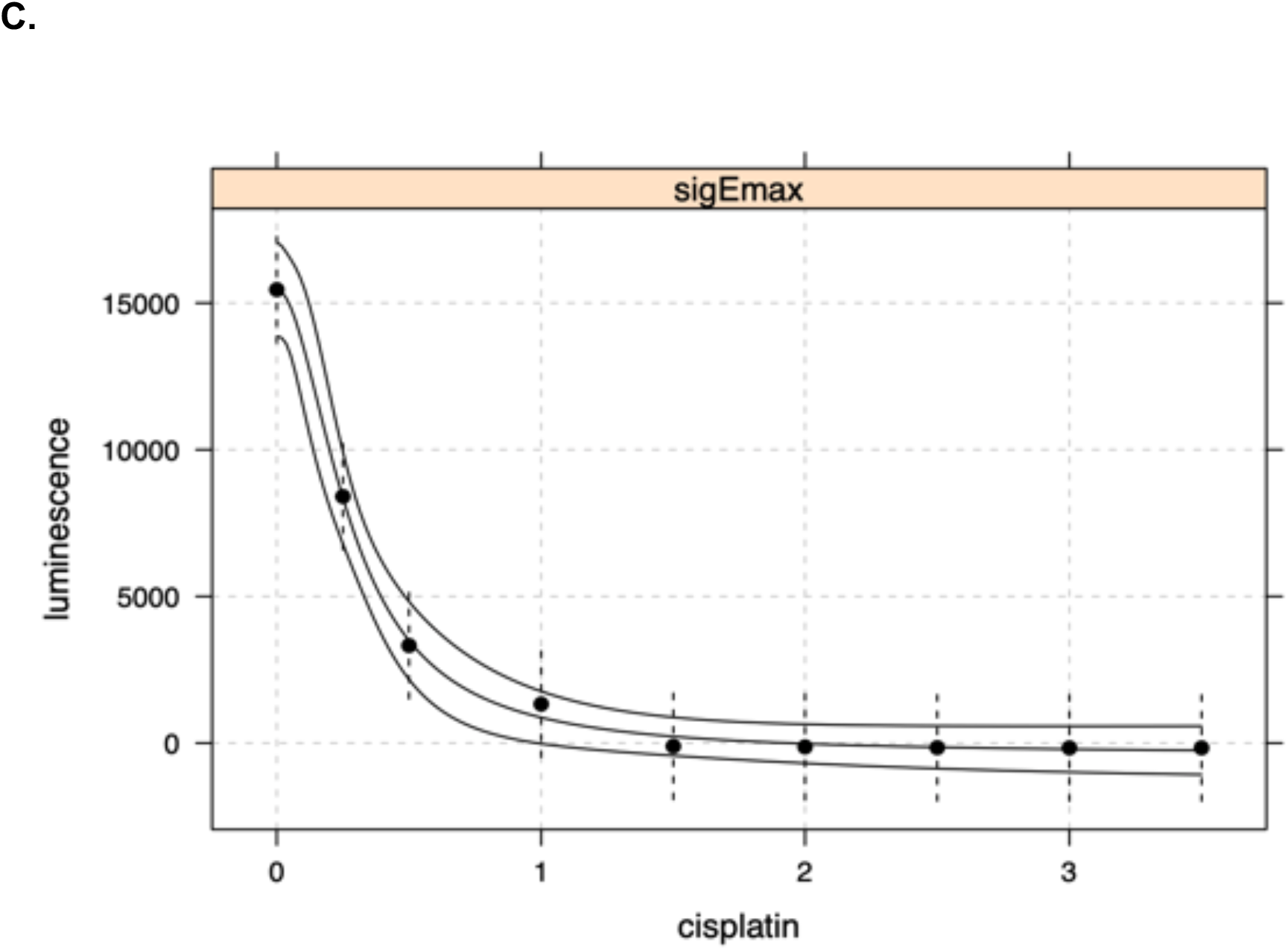
*Ex vivo* exposure-response in 2D and 3D cells incubated in Cisplatin (CIS) for 24 or 48 hours fitted with an Emax or Sigmoidal Emax Model. The X-axis shows the concentration of CIS (in μM) used and the Y-axis shows reported luminescence (proportional to the concentration of ATP in nM). Graph (A) shows results for 2D human TCC cells incubated for 24 hours in CIS. Graph (B) shows results for 2D human TCC cells incubated for 48 hours in CIS, graph (C) shows results for 3D canine TCC cells incubated for 24 hours in CIS.

**Table 1.** IC50 (potency) values obtained from the cell viability assay with Cisplatin in 2D and 3D bladder TCC cell lines.

| Figure # | Species | 2D or 3D | Drug | Exposure time | IC<sub>50</sub> [μM] |
| --- | --- | --- | --- | --- | --- |
| 6A | Human | 2D | CIS | 24-hours | 1.5 |
| 6B | Human | 2D | CIS | 48-hours | 0.25 |
| 6C | Canine | 3D | CIS | 24-hours | 0.28 |

Overall, 2D human transitional cell carcinoma cells subjected to cisplatin for 24 hours showed a 50% decrease in metabolic activity (as indicated by a 50% decrease in [ATP]) at 1.5μM of cisplatin using a simple *Imax* model (**Figure 3A**). Using the same model description (*Emax*), 2D human TCC cells subjected to Cisplatin for 48 hours showed a 50% decrease in metabolic activity at 0.25 μM of Cisplatin (**Figure 3B**), suggesting a higher potency of cisplatin after longer periods of incubation. Samples of 3D canine TCC cells subjected to cisplatin for 24 hours were better fitted with a *Sigmoidal Emax* model. The estimated IC50 value for that assay was 0.28μM of Cisplatin with a standard error of 0.03 (**Figure 3C**), so in line with IC50 estimates in 2D human TCC cell lines after 48 hours of incubation.

## DISCUSSION

In this series of experiments, we describe the culture, maintenance, and phenotypic characterization of canine, urinary-derived, TCC 3D organoids. Additionally, we subjected those TCC 3D organoids to a cell viability assay. As seen in **Figure 1**, organoids were successfully cultured and passaged. As a result of collecting organoids from 11 different canines, the opportunity to begin building a biobank arose. Organoids stored in the biobank were frozen in freezing media and stored in liquid nitrogen. Additional organoids were also paraffin-embedded in phenotypic characterization. Lastly, organoids were also stored in RNA*later™* and placed in the liquid nitrogen tank for characterization of mRNA expression. Organoids were confirmed to be of UC origin via RNAscope^®^ characterization as seen in **Figure 2**. Tissue and organoid sections from the same parent tumor were subjected to RNA *in situ* hybridization simultaneously, using the same probes. Our preliminary results showed similar levels of expression between tissues and 3D organoids for multiple RNA-based cancer cell markers (FOXA1, CD44, KRT7) (**Figure 2**). We used this information, with additional quantified RNA-ISH data not included here, to confirm the cellular identity of the organoids.

Once confirmed, we chose to run cell viability assays on 3D organoids and 2D cell lines as control. Using the standard platinum-based chemotherapeutic, cisplatin, cell viability assays were conducted on human 2D TCC cell lines and canine 3D TCC cell lines. In comparing results, 2D human UC cells exposed to cisplatin for 24 hours had an IC50 concentration lower than those exposed for 48 hours. This is expected as TCC cells exposed to cisplatin for a longer period show decreased cell viability (Wang & Wu, 2015). 3D cells treated with cisplatin for 24 hours showed the lowest IC50 values, thus suggesting that 3D organoids show increased sensitivity to cisplatin when compared to 2D cells. There could be a few factors that could explain these differences. First, the 2D and 3D cell lines were from two different species. While the 2D cells were of human origin, 3D organoids were of canine origin. Secondarily, there are differences in the extracellular environment and cell-cell interactions between 2D and 3D cells that could account for this difference. It was noted previously that certain drugs fail clinical trials as dose-response is markedly different in 2D cell culture when compared to animal models (Breslin & O’Driscoll, 2013). Canine TCC organoids performing differently than 2D cells as seen in the cell viability assay suggest that there is a notable difference between the two types of cell cultures. Next steps in this research would be to conduct additional cell viability assays on 3D cell cultures to compare to *in vivo* response in patients with bladder TCC. While more experimenting is necessary, via genomic characterization and more cell viability assays spanning longer time points (48 hours, 72 hours), 3D cells canine TCC cell culture should be explored as the next potential model to study bladder TCC *ex vivo*.

## Footnotes

1 R Core Team (2020). R: A language and environment for statistical computing. R Foundation for Statistical Computing, Vienna, Austria. URL https://www.R-project.org/.

## REFERENCES

Ploeg M, Aben KKH, Kiemeney LA. The present and future burden of urinary bladder cancer in the world. World J Urol. 2009 Jun;27(3):289–93.

Siegel RL, Miller KD, Jemal A. Cancer statistics, 2020. CA Cancer J Clin. 2020;70(1):7–30.

Park JC, Citrin DE, Agarwal PK, Apolo AB. Multimodal management of muscle invasive bladder cancer. Curr Probl Cancer. 2014;38(3):80–108.

Abraham J, Gulley JL, Allegra CJ. The Bethesda Handbook of Clinical Oncology. Lippincott Williams & Wilkins; 2012. 682 p.

Kim LHC, Patel MI. Transurethral resection of bladder tumour (TURBT). Transl Androl Urol. 2020 Dec;9(6):3056072–3053072.

Aragon-Ching JB, Werntz RP, Zietman AL, Steinberg GD. Multidisciplinary Management of Muscle-Invasive Bladder Cancer: Current Challenges and Future Directions. Am Soc Clin Oncol Educ Book. 2018 May 23;(38):307–18.

Stein JP, Lieskovsky G, Cote R, Groshen S, Feng A-C, Boyd S, et al. Radical Cystectomy in the Treatment of Invasive Bladder Cancer: Long-Term Results in 1,054 Patients. J Clin Oncol. 2001 Feb 1;19(3):666–75.

Thomsen MBH, Nordentoft I, Lamy P, Høyer S, Vang S, Hedegaard J, et al. Spatial and temporal clonal evolution during development of metastatic urothelial carcinoma. Mol Oncol. 2016;10(9):1450–60.

Thomsen MBH, Nordentoft I, Lamy P, Vang S, Reinert L, Mapendano CK, et al. Comprehensive multiregional analysis of molecular heterogeneity in bladder cancer. Sci Rep [Internet]. 2017 Sep 15.

Edmondson R, Broglie JJ, Adcock AF, Yang L. Three-dimensional cell culture systems and their applications in drug discovery and cell-based biosensors. Assay Drug Dev Technol. 2014 May;12(4):207–18.

Birgersdotter A, Sandberg R, Ernberg I. Gene expression perturbation ex vivo--a growing case for three-dimensional (3-D) culture systems. Semin Cancer Biol. 2005 Oct;15(5):405–12.

Hutchinson L, Kirk R. High drug attrition rates--where are we going wrong? Nat Rev Clin Oncol. 2011 Mar 30;8(4):189–190. PMID: 21448176.

Wittenburg LA, Gustafson DL. Optimizing preclinical study design in oncology research. Chemico-Biological Interactions [Internet]. 2011 Apr [cited 2021 Feb 11];190(2–3):73–78.

Kumar S, Bajaj S, Bodla R. Preclinical screening methods in cancer. Indian J Pharmacol [Internet]. 2016 [cited 2021 Feb 11];48(5):481.

Nixon NA, Khan OF, Imam H, Tang PA, Monzon J, Li H, Sun G, Ezeife D, Parimi S, Dowden S, Tam VC. Drug development for breast, colorectal, and non-small cell lung cancers from 1979 to 2014: Cancer Drug Development From 1979-2014. Cancer [Internet]. 2017 Dec 1 [cited 2021 Feb 11];123(23):4672–4679.

Ruan J-L, Hsu J-W, Browning RJ, Stride E, Yildiz YO, Vojnovic B, Kiltie AE. Mouse Models of Muscle invasive Bladder Cancer: Key Considerations for Clinical Translation Based on Molecular Subtypes. Eur Urol Oncol. 2019 May;2(3):239–247. PMID: 31200837.

Arantes-Rodrigues R, Colaço A, Pinto-Leite R, Oliveira PA. Ex vivo and In Vivo Experimental Models as Tools to Investigate the Efficacy of Antineoplastic Drugs on Urinary Bladder Cancer. Anticancer Res. 2013 Apr 1;33(4):1273–96.

Kobayashi T, Owczarek TB, McKiernan JM, Abate-Shen C. Modeling bladder cancer in mice: opportunities and challenges. Nat Rev Cancer. 2015 Jan;15(1):42–54.

Knapp DW, Ramos-Vara JA, Moore GE, Dhawan D, Bonney PL, Young KE. Urinary Bladder Cancer in Dogs, a Naturally Occurring Model for Cancer Biology and Drug Development. ILAR J. 2014 Jan 1;55(1):100–18.

Vasyutin I, Zerihun L, Ivan C, Atala A. Bladder Organoids and Spheroids: Potential Tools for Normal and Diseased Tissue Modelling. Anticancer Res. 2019 Mar 1;39(3):1105–18.

Fatehullah A, Tan SH, Barker N. Organoids as an ex vivo model of human development and disease. Nat Cell Biol. 2016 Mar;18(3):246–54.

Hughes CS, Postovit LM, Lajoie GA. Matrigel: A complex protein mixture required for optimal growth of cell culture. PROTEOMICS. 2010;10(9):1886–90.

Mochel JP, Jergens AE, Kingsbury D, Kim HJ, Martín MG, Allenspach K. Intestinal Stem Cells to Advance Drug Development, Precision, and Regenerative Medicine: A Paradigm Shift in Translational Research. AAPS J. 2017 Dec 12;20(1):17. DOI: 10.1208/s12248-017-0178-1. Review. PMID: 29234895.

Chandra L, Borcherding DC, Kingsbury D, Atherly T, Ambrosini YM, Bourgois-Mochel A, Yuan W, Kimber M, Qi Y, Wang Q, Wannemuehler M, Ellinwood NM, Snella E, Martin M, Skala M, Meyerholz D, Estes M, Fernandez-Zapico M, Mochel JP* and Allenspach K*. Derivation of Adult Canine Intestinal Organoids for Translational Research in Gastroenterology. BMC Biol. 2019 Apr 11;17(1):33. DOI: 10.1186/s12915-019-0652-6. PMID: 30975131.* co-corresponding author.

Ambrosini YM, Park Y, Jergens AE, Shin W, Mon S, Atherly T, Borcherding DC, Jang J, Allenspach K, Mochel JP* and Kim HY*. Recapitulation of an Accessible Interface of the Biopsy-Derived Canine Intestinal Organoids to Study Epithelial-Luminal Interactions. PLoS One. 2020 Apr 17;15(4):e0231423. DOI: 10.1371/journal.pone.0231423. PMID: 32302323. *: co-corresponding author.

Fulkerson CM, Dhawan D, Ratliff TL, Hahn NM, Knapp DW. Naturally Occurring Canine Invasive Urinary Bladder Cancer: A Complementary Animal Model to Improve the Success Rate in Human Clinical Trials of New Cancer Drugs. Int J Genomics. 2017 Apr 9;2017:e6589529.

Minkler S, Lucien F, Kimber MJ, Sahoo DK, Bourgois-Mochel A, Musser M, Johannes C, Frank I, Cheville J, Allenspach K, Mochel JP. Emerging Roles of Urine-Derived Components for the Management of Bladder Cancer: One Man’s Trash Is Another Man’s Treasure. Cancers (Basel). 2021 Jan 23;13(3):422. DOI: 10.3390/cancers13030422. PMID: 33498666.

Iennarella-Servantez CA, Gabriel V, Atherly T, (…), Bourgois-Mochel A, Jergens AE, Allenspach K, Mochel JP. Collection, Culture, and Characterization of Canine Healthy Bladder and Urothelial Carcinoma Organoids: Reverse Translational Clinical Research in the Veterinary Patient. 2021. European College of Veterinary Internal Medicine Annual Conference (Virtual).

Gabriel V, Iennarella-Servantez CA, Atherly T, Minkler S, Thenuwara S, Mao S, Colosimo M, Kurr L, Borcherding D, Bourgois-Mochel A, Jergens AE, Mochel JP, Allenspach K. Culture and Maintenance of Well-Differentiated Canine Hepatic Organoids and Urinary Bladder Organoids. 2021. European College of Veterinary Internal Medicine Annual Conference (Virtual).

Valli VE, Norris A, Jacobs RM, Laing E, Withrow S, Macy D, et al. Pathology of canine bladder and urethral cancer and correlation with tumour progression and survival. J Comp Pathol. 1995 Aug 1;113(2):113–30.

Knapp DW, Dhawan D, Ostrander E. “Lassie,” “Toto,” and Fellow Pet Dogs: Poised to Lead the Way for Advances in Cancer Prevention. Am Soc Clin Oncol Educ Book. 2015 May 1;(35):e667–72.

Knapp DW, Glickman NW, Widmer WR, DeNicola DB, Adams LG, Kuczek T, Bonney PL, DeGortari AE, Han C, Glickman LT. Cisplatin versus cisplatin combined with piroxicam in a canine model of human invasive urinary bladder cancer. Cancer Chemother Pharmacol. 2000;46(3):221–6. doi: 10.1007/s002800000147. PMID: 11021739.

Boria PA, Glickman NW, Schmidt BR, Widmer WR, Mutsaers AJ, Adams LG, Snyder PW, DiBernardi L, de Gortari AE, Bonney PL, Knapp DW. Carboplatin and piroxicam therapy in 31 dogs with transitional cell carcinoma of the urinary bladder. Vet Comp Oncol. 2005 Jun;3(2):73–80. doi: 10.1111/j.1476-5810.2005.00070.x. PMID: 19379215.

Robat C, Burton J, Thamm D, Vail D. Retrospective evaluation of doxorubicin-piroxicam combination for the treatment of transitional cell carcinoma in dogs. J Small Anim Pract. 2013 Feb;54(2):67–74. doi: 10.1111/jsap.12009. Epub 2013 Jan 3. PMID: 23286739.

Ambrosini YM, Park Y, Jergens AE, Shin W, Min S, Atherly T, et al. Recapitulation of the accessible interface of biopsy-derived canine intestinal organoids to study epithelial-luminal interactions. PLOS ONE. 2020 Apr 17;15(4):e0231423.

Wang F, Flanagan J, Su N, Wang L-C, Bui S, Nielson A, et al. RNAscope. J Mol Diagn JMD. 2012 Jan;14(1):22–9.

Gerhardt RT, Perras JP, Sevin BU, Petru E, Ramos R, Guerra L, et al. Characterization of ex vivo chemosensitivity of perioperative human ovarian malignancies by adenosine triphosphate chemosensitivity assay. Am J Obstet Gynecol. 1991 Aug;165(2):245–55.

DeGraff DJ, Clark PE, Cates JM, Yamashita H, Robinson VL, Yu X, et al. Loss of the Urothelial Differentiation Marker FOXA1 Is Associated with High Grade, Late Stage Bladder Cancer and Increased Tumor Proliferation. PLoS ONE [Internet]. 2012 May 10 [cited 2021 Apr 24];7(5).

Osei-Amponsa V, Buckwalter JM, Shuman L, Zheng Z, Yamashita H, Walter V, et al. Hypermethylation of FOXA1 and allelic loss of PTEN drive squamous differentiation and promote heterogeneity in bladder cancer. Oncogene. 2020 Feb;39(6):1302–17.

Wu C-T, Lin W-Y, Chang Y-H, Chen W-C, Chen M-F. Impact of CD44 expression on radiation response for bladder cancer. J Cancer. 2017 Apr 9;8(7):1137–44.

Ichimi T, Enokida H, Okuno Y, Kunimoto R, Chiyomaru T, Kawamoto K, et al. Identification of novel microRNA targets based on microRNA signatures in bladder cancer. Int J Cancer. 2009;125(2):345–52.

Wang D, Wu X. *In vitro* and *in vivo* targeting of bladder carcinoma with metformin in combination with cisplatin. Oncol Lett. 2015 Aug;10(2):975–981.

Breslin S, O’Driscoll L. Three-dimensional cell culture: the missing link in drug discovery. Drug Discov Today. 2013 Mar;18(5–6):240–9.

